# Dissociating harmonic and non-harmonic phase-amplitude coupling in the human brain

**DOI:** 10.1101/2020.08.11.246298

**Authors:** Janet Giehl, Nima Noury, Markus Siegel

## Abstract

Phase-amplitude coupling (PAC) has been hypothesized to coordinate cross-frequency interactions of neuronal activity in the brain. However, little is known about the distribution of PAC across the human brain and the frequencies involved. Furthermore, it remains unclear to what extend PAC may reflect spurious cross-frequency coupling induced by physiological artifacts or rhythmic non-sinusoidal signals with higher harmonics. Here, we combined MEG, source-reconstruction and different measures of cross-frequency coupling to systematically characterize PAC across the resting human brain. We show that cross-frequency measures of phase-amplitude, phase-phase, and amplitude-amplitude coupling are all sensitive to signals with higher harmonics. In conjunction, these measures allow to distinguish harmonic and non-harmonic PAC. Based on these insights, we found no evidence for non-harmonic PAC in resting-state MEG. Instead, we found cortically and spectrally wide-spread PAC driven by harmonic signals. Furthermore, we show how physiological artifacts and spectral leakage cause spurious PAC across wide frequency ranges. Our result clarify how different measures of cross-frequency interactions can be combined to characterize PAC and cast doubt on the presence of prominent non-harmonic phase-amplitude coupling in human resting-state MEG.

## 1. Introduction

Oscillations of neuronal activity are abundant in the brain and have been implicated in various cognitive processes (Siegel et al., 2012). Recently, the question how different oscillations and their associated networks interact has received considerable attention (Lisman and Idiart, 1995; Jensen and Colgin, 2007; Canolty and Knight, 2010; Tort et al., 2010; Jensen et al., 2012; Aru et al., 2015; Colgin, 2015; Hyafil et al., 2015; McLelland and VanRullen, 2016; Dvorak and Fenton, 2014; Fell and Axmacher, 2011). The most prominently discussed mode of cross-frequency coordination has been phase-amplitude coupling (PAC), where the phase of a neuronal oscillation is correlated with the amplitude of another comparatively faster neuronal oscillation (Canolty et al., 2006). It has been suggested that PAC could not only link oscillations within one area, but could also play a role in long-range interactions between areas (Jensen and Colgin, 2007; Siebenhühner et al., 2020; van der Meij et al., 2012; von Nicolai et al., 2014; Nandi et al., 2019).

However, so far PAC has not been systematically mapped across the human brain and across frequency combinations. Furthermore, the interpretation of cross-frequency coupling results is challenging. The biggest caveat is that measures of PAC are not only sensitive to the phase-amplitude coupling between two different oscillations of interest, but are also sensitive to signals with higher harmonics, i.e. signals that contain a periodic, but non-sinusoidal component. Therefore, there is an inherent ambiguity in the results from these measures. In fact, several studies have cautioned for or reported that harmonic PAC is linked to non-sinusoidal waveform shapes of oscillations rather than the non-harmonic PAC of interest (Aru et al., 2015; Chacko et al., 2018; Cole et al., 2017; Cole and Voytek, 2017; Gerber et al., 2016; Hyafil, 2015; Jensen et al., 2016; Kramer et al., 2008; Lozano-Soldevilla et al., 2016; Vaz et al., 2017; Velarde et al., 2019). Therefore, it is challenging to not only map, but to also correctly characterize the nature of any measured cross-frequency coupling.

Here, we addressed this question, by evaluating phase-amplitude coupling across a broad range of frequency combinations and the entire human cortex based on two independent source-reconstructed magnetoencephalography (MEG) datasets. We estimated PAC using two different measures: vectorlength-PAC (Canolty et al., 2006) and bicoherence (Shahbazi Avarvand et al. 2018). To test if the measured PAC patterns reflect non-harmonic neuronal PAC, we assessed, first, if these patterns reflect muscle- or eye-movement artifacts, and second, to what extent they are affected by the non-sinusoidal shape of neuronal oscillations. To this end, we systematically applied a novel procedure that allows to distinguish harmonic and non-harmonic PAC.

## 2. Materials and Methods

### 2.1. Datasets

We used resting-state MEG data from two different datasets: the Human Connectome Project MEG data (HCP) (Van Essen et al., 2013), which is publicly available, and an independent MEG dataset recorded at the MEG-Center Tübingen.

### 2.2. Data acquisition

Unless specified otherwise, set-up, recording, and preprocessing of the HCP dataset (HCP S1200 Release), was as previously described (Van Essen et al., 2013). We used the first of three sessions of 6 minutes eyes-open resting-state MEG, which was available for 89 subjects. Subjects were in supine position and fixated a red fixation cross on dark background.

The Tübingen dataset comprised 28 healthy subjects (17 female, mean age 26.5 years), that all gave written informed consent and received monetary compensation. For this dataset, 10 minutes of resting-state MEG were recorded with 275 channels at a sampling rate of 2,343.75 Hz (Omega 2000, CTF Systems, Inc., Port Coquitlam, Canada). Participants were seated upright in a dimly lit magnetically shielded chamber and fixated a central fixation point. The recordings was approved by the local ethics committee and conducted in accordance to the Declaration of Helsinki. All participants gave written informed consent before participating. For both datasets, structural T1-weighted MRIs of all subjects were used to construct individual head and source models.

### 2.3. Preprocessing - HCP dataset

We down sampled the data to 1000 Hz, band-pass filtered between 0.1 and 400 Hz, and notch-filtered between 59 and 61 Hz (and harmonics) using zero-phase 4^th^-order Butterworth forward and reverse filters. We removed artifactual data segments as defined by the HCP pipeline (baddata). We manually identified and removed muscle-, eye- and heart-related artifacts using ICA (Hipp and Siegel, 2013). Heart-related artifact ICs were removed in all 89 subjects. Eye-related ICs were removed in 86 subjects. In this dataset, muscle-related ICs often contained prominent signal components in the alpha-band. To avoid accidentally removing components that may include neuronal interactions, we did not remove these components. Thus, we removed muscle-ICs in only 30 of the 89 subjects, which resulted in considerable residual muscle activity in this preprocessed dataset.

### 2.4. Preprocessing - Tübingen dataset

The data was down sampled to 1000 Hz, low-pass filtered at 300 Hz, notch-filtered between 49 and 51 Hz (and harmonics) and high-pass filtered at 0.5 Hz using zero-phase 4^th^-order Butterworth forward and reverse filters. Segments with jumps, eye blinks or strong muscle activity were removed manually in the time domain data. Remaining artifacts were removed using ICA analysis as described above for the HCP dataset. For this dataset, however, we removed muscle-ICs more stringently even if they contained a spectral peak in the alpha-band. Thus, for this dataset muscle-related ICs were removed in all subjects.

### 2.5. Source reconstruction

For both datasets, we used beamforming to reconstruct cortical activity at 457 positions on a shell that covered the entire cortex with even spacing approximately 1 cm beneath the skull (Hipp and Siegel, 2015). For the HCP dataset, we used the spatial transformation matrices that are available in the individual source models provided with the HCP dataset and applied them to our source model. We performed two distinct source space projections and frequency analyses for vectorlength-PAC and all other cross-frequency measures.

### 2.6. Spectral analysis and beamforming for vectorlength-PAC

We employed Morlet wavelets with logarithmically scaled center frequencies ranging from 2^−0.25^ Hz (~0.84 Hz) to 2^8^ Hz (256 Hz) in quarter-octave steps (factor of 2^−0.25^).

Optimal for PAC detection is a relatively low frequency resolution for amplitude frequencies that must include the amplitude modulation side-peaks and a relatively high frequency resolution for the phase frequency (Aru et al., 2015). Therefore, we chose bandwidths of 0.5 octaves for the phase frequencies and 1 octave for the amplitude frequencies (full width at half maximum). This corresponds to a wavelet-width *q* = *f*/*σ*_*f*_ ~6.86 for 0.5 octave and *q* = *f*/*σ*_*f*_ ~3.53 for 1 octave. We computed time-frequency estimates with a temporal steps size of 7ms. Subsequently, we applied DICS beamforming with frequency specific filters (Gross et al., 2001) For each source, we computed filters in the dominant dipole direction.

### 2.7. Vectorlength-PAC

We estimated vectorlength phase-amplitude coupling between phase-frequencies (*f*_*φ*_) of ~0.84 Hz (2^−0.25;^ Hz) to 128 Hz (2^7^ Hz) and amplitude-frequencies (*f*_*A*_) of at least twice the phase frequency. We estimated vectorlength-PAC *V(f)* according to (Canolty et al., 2006):

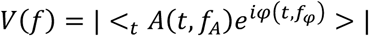

Here, *φ*(*t*, *f*_*φ*_) is the phase, *A* (*t*, *f*_*A*_) the amplitude, <_*t*_,…> is the average over time and |…| is the absolute value.

### 2.8. Spectral analysis and beamforming for other cross-frequency measures

For all other cross-frequency coupling measures (bicoherence, cross-frequency amplitude-amplitude correlation, cross-frequency phase-phase coupling) we performed linearly constrained minimum variance (LCMV) beamforming (Van Veen et al., 1997) to estimate source level activity. This approach preserves the phase relationship between frequencies. The source-data was split into half-overlapping 1 s segments (2 s and 4s segments for two control analyses related to spectral leakage). Each segment was demeaned and a Hanning window was applied. Then, fast Fourier transformation was computed for each segment using zero-padding to 2 or 10 seconds, which results in frequency resolutions of 0.5 or 0.1 Hz, respectively.

### 2.9. Bicoherence

We estimated Bicoherence for frequencies *f*_1_ (0.5 to 64 Hz) and *f*_2_ (1 to 200 Hz) in steps of 0.5 Hz with *f*_2_ ≥ *f*_1_ − 3 Hz and the corresponding *f*_2_ = *f*_1_ + *f*_2_ according to the formula with the normalization factor from (Hagihira et al., 2001):

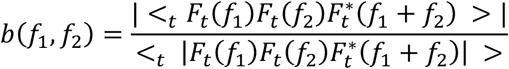

Here, *F*_*t*_(*f*) is the signal’s time-frequency transformation at time t, | … | represents the absolute value, and <_*t*_ … > is the average over time. We use the terms “*f*_2_” and “*f*_1_ + *f*_2_” interchangeably throughout the paper. Frequencies *f*_2_ smaller than *f*_1_ were included for accurate detection of bicoherence peaks at and close to *f*_1_ = *f*_2_, in particular after smoothing of the cross-frequency spectrum.

To locate delta-*f*_1_-range bicoherence peaks, bicoherence was estimated in steps of 0.1 Hz for *f*_1_ (0.1 to 2.5 Hz) and 0.5 Hz for *f*_2_ (0.5 to 50 Hz), with *f*_2_ ≥ *f*_1_.

### 2.10. Cross-frequency amplitude-amplitude coupling (AAC)

We computed the Pearson correlation coefficients between amplitude time-series, that were derived from the LCMV-beamformed Fourier time-series, between frequencies *f*_1_ (0.5 to 64 Hz) and *f*_2_ (1 to 200 Hz) in steps of 0.5 Hz where *f*_2_ ≥ *f*_1_ + 3 Hz.

### 2.11. Cross-frequency phase-phase coupling (PPC)

We computed cross-frequency phase-coupling between frequencies *n* and *m* weighted and normalized by the signal amplitudes at these frequencies. The phase of the lower frequency *n* was accelerated by the factor *m*/*n* to match the higher frequency *m*. The resulting measure can be understood as computing coherence between signals of different frequencies.

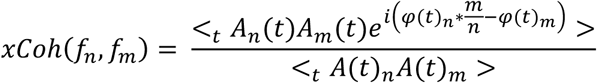

<_*t*_ … > represents the average over time. Amplitude *A* (*t*) and phase estimates *φ*(*t*) were obtained from the time-frequency Fourier estimates. The measure approaches 0 for a random phase relationship and equals 1 for perfect m/n factor phase-coupling. The normalization matches the normalization used for Bicoherence. Furthermore, the weighting by the signal amplitudes improves the robustness of the measure by favoring phase information during times of high signal amplitude. We computed this measure for the same frequency pairs as AAC.

### 2.12. Regions of interest (ROIs)

We defined 4 ROIs based on the local maxima of the group-level vectorlength-PAC averaged across frequencies in left/right sensorimotor (MNI: −44.4 −30.2 58.9 / 39.7 −20.0 61.0), visual (−19.1 −99.7 2.1 / 19.1 −99.6 1.7), parietal (−11.1 −81.3 45.3 / 11.1 −81.6 45.4) and superior temporal cortex (−62.5 −43.9 15.0 / 63.0 −8.6 15.0). We averaged coupling measures across hemispheres.

### 2.13. Statistical analysis of coupling measures

We employed a time-shifting surrogate procedure to assess the statistical significance of cross-frequency coupling. We generated 100 random circular time-shifts. The same time-shift was applied for all sources and frequency combinations. For the vectorlength measure, the amplitude time-series of each subject was shifted circularly relative to the phase time-series and vectorlength-PAC was computed for each of the 100 shifts. For bicoherence, we shifted the time-series of *f*_3_ relative to *f*_1_ and *f*_2_. For AAC and PPC, we shifted the timeseries of *f*_2_ relative to *f*_1_. AAC values and corresponding surrogate values were Fisher-z-transformed. For each measured coupling measure, a z-score was computed relative to the corresponding 100 surrogate values. To determine significant cross-frequency coupling while controlling for multiple comparisons, we employed a cluster permutation statistic across cross-frequency space or cortical space. For bicoherence and PPC statistics, cross-frequency spectra were smoothed with a 5-by-5 frequency-bin sized Hanning window. Clusters across cortical and cross-frequency space were defined based on the t-statistic of cross-frequency measures across subjects with a cluster threshold of p < 0.01. Clusters were then defined under the null-hypothesis by random sign flip across subjects before computing the t-statistic (1000 repeats). Cluster significance was defined as the probability to obtain a maximum cluster of at least this size across the null-hypothesis permutations with p < 0.05.

### 2.14. Analysis of the influence of artifacts

To study the potential influence of residual artifacts on the cross-frequency coupling results, we compared the spectral and spatial PAC patterns of artifactual signals defined during data preprocessing to the results of the cleaned data (Hipp and Siegel, 2013). To this end, we back-projected only the artifactual ICs to the sensors, beamformed these artifactual signals, and computed vectorlength-PAC and bicoherence on the source-localized artifacts. Finally, we compared the patterns of the artifactual signals with the patterns of the cleaned signal. As residual heart-artifacts tend to be project to the brain center, we restricted this analysis to muscle- and eye-related artifacts.

### 2.15. Simulated signals with harmonic and non-harmonic PAC

We computed bicoherence, AAC and PPC of simulated signals with either higher harmonics or non-harmonic phase-amplitude coupling. These simulations illustrate typical cross-frequency coupling patterns expected for different signal types and cross-frequency measures.

In one of the non-harmonic PAC signals, the carrier frequency (*f*_*c*_) and modulation side-peaks were distinct from the modulating frequency (*f*_*m*_). To this end, we simulated coupling between *f*_*m*_=10 Hz and *f*_*c*_=40 Hz (resulting in side-peaks at 30 Hz and 50 Hz) with 10-times smaller amplitude for *f*_*c*_ than *f*_*m*_. Additionally, we simulated the special case of PAC, where the lower side-peak of the amplitude modulation (*f*_*c-m*_) coincides with the modulating frequency (*f*_*m*_=10 Hz and *f*_*c*_=20 Hz, resulting in side-peaks at 10 Hz and 30 Hz, and with 4 times smaller amplitude for *f*_*c*_ than *f*_*m*_).

For obtaining the raw signals at *f*_*m*_ and *f*_*c*_ (*x*_*fm*_(*t*) and *x*_*fc*_(*t*), respectively), we took 6 minutes of white noise and, at each time point, applied a Hanning window, calculated the FFT, extracted the frequency bin of interest, and applied the inverse Fourier transform. For each frequency, a Hanning window with a defined spectral width (full width at half maximum, FWHM) was used (FWHM = 1.5 Hz for *f*_*m*_, and FWHM = 3 Hz and 2 Hz for *f*_*c*_, for the two different signals, respectively). Then, non-harmonic PAC signals were generated by amplitude modulation of *x*_*fc*_ according to the phase of *x*_*fm*_ as follows:

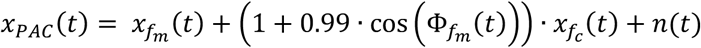

Φ_*fm*_(*t*) is the phase of *x*_*fm*_, extracted from the above mentioned time-Fourier analysis. *n* (*t*) is 1/*f*^2^-noise.

For the harmonic-PAC signal, we obtained the base component at 10 Hz (*x*_*f10*_(*t*)) using the procedure explained above for *f*_*c*_, and added it up with 4 higher harmonics and 1/*f*^2^-noise (*n* (*t*)). Higher harmonics of *x*_*f10*_ were generated as:

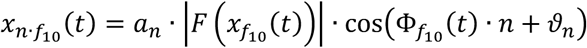

Here, |*F* (*x*_*f10*_(*t*)| and Φ_*f10*_(*t*) are the instantaneous amplitude- and phase-time series of *x*_*f10*_, respectively, and n is the integer multiplier (*n* ∈ {2,3,4,5}). The relative amplitude and phase of the higher harmonics relative to the base component were controlled by the factors *a*_*n*_ and *θ*_*n*_, respectively (*a*_2_=0.25, *a*_3_=0.06, *a*_4_=0.025, *a*_5_=0.05, *ϑ*_2_=pi/2, *ϑ*_3_=pi/4, *ϑ*_4_=3/2*pi, and *ϑ*_5_=2*pi).

For all simulated signals, we applied the same processing as for the measured source-level data and calculated bicoherence, cross-frequency AAC and PPC as well as a time-average log-power spectrum.

### 2.16. Peak localization

To localize peaks in the frequency-frequency space for bicoherence, AAC, and PPC, we averaged the cross-frequency values across *f*_2_ frequencies below 70 Hz. Then, we identified the *f*_1_ with the maximum averaged cross-frequency measure (in the range 5 to 14 Hz for single subjects, 7.5 to 14 Hz for the average across subjects, and 0.1 to 2.5 Hz for the delta-*f*_1_ peaks). Finally, we identified the corresponding *f*_2_ frequency maxima for each maximum *f*_1_.

### 2.17. Single-subject harmonic peaks in bicoherence

After observing harmonic peaks of theta/alpha PAC averaged across subjects, we tested for such a harmonic pattern across subjects. Specifically, we tested if *f*_2_ frequencies of bicoherence peaks were at integer multiples of corresponding *f*_1_ frequencies.

For localizing bicoherence peaks, we first smoothed the Z-scored bicoherence cross-frequency spectra by convolution with a 3×3 Hanning window. To more precisely detect peaks close to the *f*_1_ = *f*_2_ diagonal of the bicoherence cross-frequency spectrum, we extended the lower range of *f*_2_-frequencies to *f*_1_-3 Hz, but not lower than 0.5 Hz. Then, we created individual masks of single-subject significance by converting the Z-scores to p-values, false-discovery-rate correcting the p-values for multiple comparisons (Benjamini and Hochberg, 1995), and creating masks at the alpha level of 0.05. Next, we spline-interpolated both, the Z-scored cross-frequency spectra and masks, to 0.1 Hz resolution. Finally, we identified peaks of significant bicoherence as detailed above.

If bicoherence peaks reflected harmonics of an oscillator at the corresponding *f*_1_ frequency, they should fit the following regression model:

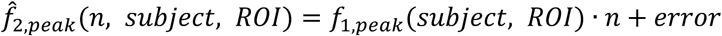

Here, *n* refers to the peak number. To avoid mislabeling of harmonic peaks, e.g. when a lower peak was not detectable, we set *n* to match the closest harmonic. In total, up to 6% of peaks per ROI were relabeled. For two peaks falling closest to the same harmonic, we omitted the peak associated with the weaker bicoherence Z-score. To rule out any bias due to peak relabeling, we tested the model fit against a permutation statistic that also entailed peak relabeling. For each ROI, we took peak positions of all subjects and 1000 times randomly reassigned *f*_2_ and *f*_1_ values across subjects, with the restriction that the same number of peaks had to fall within the 1^st^, 2^nd^, etc. peak groups as observed in the original data. For each of the 1000 surrogates, as well as for the original data, we computed the coefficient of determination (*R*^2^) as a measure of model fit for each regression model (one for each n^th^ harmonic at each ROI). Finally, we obtained p-values by comparing the original *R*^2^ against the surrogates.

### 2.18. Simulating the Bicoherence leakage pattern

In order to identify conspicuous caveats in the bicoherence analysis, such as leakage into remote frequency ranges, we simulated a 10-minute long signal (1000 Hz sampling rate) with higher harmonics that resembled the motor mu-rhythm with little noise and computed its Bicoherence the same way as we did for a source reconstructed signal. To construct this signal, we took the same steps as explained above, with the only difference that the base component was reconstructed as *x*_*f10*_(*t*) = 1 · cos(*φ*_10_(*t*)). The applied Hanning window was 4-s long, and the final signal was constructed using the following parameters: *a*_2_=0.35, *a*_3_=0.2, *a*_4_=0.05, *ϑ*_2_=pi, *ϑ*_3_=0, and *ϑ*_4_=pi.

### 2.19. Bicoherence leakage pattern

To test the observation that some of the Bicoherence peaks with delta-*f*_1_ reflect the leakage pattern of noisy harmonic signals of the alpha range, we made use of the predictable location of the leakage pattern. We constructed regression models to predict peak positions and tested the model fit of the data against a permutation statistic. Specifically, we applied the following regression model:

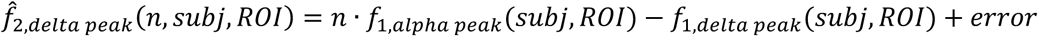

Here, for each subject and ROI, *f*_1,delta peak_ represents delta range *f*_1_ of the peaks, 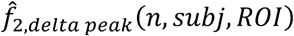 represents the *n*-th predicted *f*_2_ peak corresponding to *f*_1,delta peak_, and *f*_1,alpha peak_ represents the alpha-range *f*_1_-frequency that corresponds to the Bicoherence peaks in that range. For each subject and ROI, *f*_1, alpha peak_ was estimated as described above.

To locate (*f*_1,delta peak_, *f*_2,delta peak_) pairs more precisely, we used the z-scored bicoherence estimates with step size of 0.1 Hz for *f*_1_ frequencies and 0.5 Hz for *f*_2_ frequencies. Next, we interpolated the cross-frequency spectra along their *f*_2_ axis to steps of 0.1 Hz, and smoothed the results by convolving with a 3×3 Hanning kernel. Then, we localized the peaks at delta-range *f*_1_ frequencies.

### 2.20. Analysis software

All analyses were performed in MATLAB (MathWorks Inc., Natick, USA) using the Fieldtrip toolbox (Oostenveld et al., 2011) and custom software.

## 3. Results

### 3.1. Spectral structure of phase-amplitude coupling

We estimated vectorlength-PAC (Canolty et al., 2006) between a wide range of frequency pairs for 457 cortical sources in 89 subjects of the Human Connectome Project (HCP) resting-state MEG dataset (Fig. 1A). Averaged across the entire cortex, we observed significant coupling peaks in three different frequency ranges (Fig. 1A, top): between gamma phase frequencies and high gamma amplitude frequencies peaking at [*f*_*φ*_, *f*_*A*_] = [76.1Hz, 152.2Hz], between alpha phase frequencies and beta amplitude frequencies peaking at [*f*_*φ*_, *f*_*A*_] = [11.3Hz, 22.6Hz], and weak but significant between delta phase-frequencies and amplitude frequencies around 16 Hz. In the cortical space and averaged over all frequency pairs (Fig. 1A, bottom), the vectorlength measure peaked primarily in bilateral sensorimotor regions, as well as in temporal, occipital and parietal areas. Do these patterns reflect true neuronal interactions between distinct oscillatory processes? To answer this, we first tested if these patterns were related to non-neuronal signals, i.e. muscle- or eye-movement artifacts.

**Figure 1.**
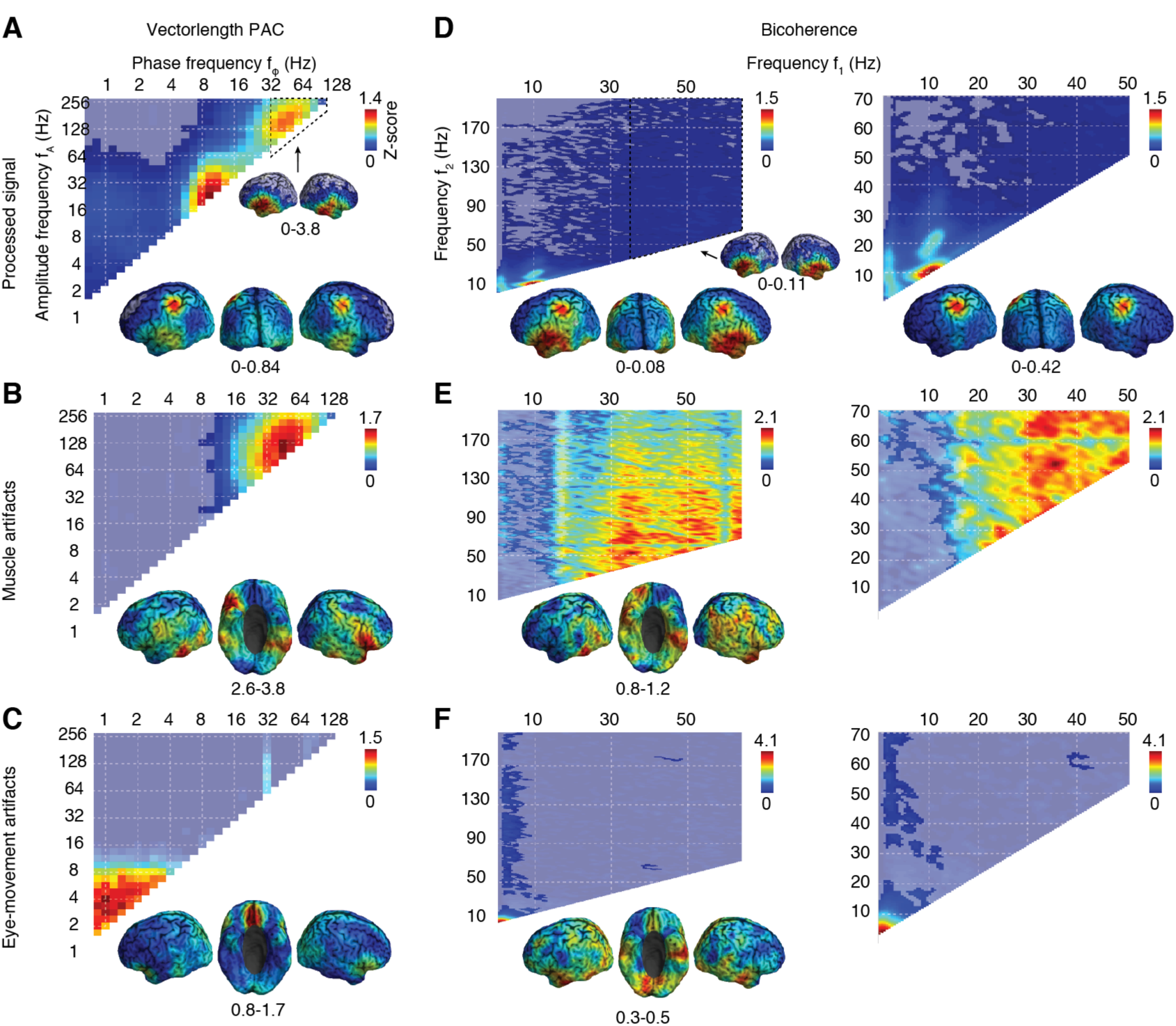
Vectorlength-PAC and bicoherence. (**A**) Vectorlength-PAC averaged across the cortex (top), and its cortical distribution, averaged either across all (bottom) or high gamma frequencies (inset). (**B**) Vectorlength-PAC of rejected muscle artifacts (independent components) averaged across the cortex (top) and all significant frequency combinations (bottom). (**C**) Vectorlength-PAC of rejected eye-related artifacts (independent components) averaged across the cortex (top) and all significant frequency combinations (bottom). (**D**) Bicoherence averaged across the cortex for a large frequency range (left) and zoomed-in lower frequency range (right). Cortical distributions of Bicoherence averaged for the corresponding frequency ranges are shown below. The inset shows the cortical distribution of bicoherence for a high-frequency range often associated with muscle activity. (**E**) Bicoherence of rejected muscle artifacts averaged across the cortex (top) and for significant frequency combinations (bottom) (**F**) Bicoherence of rejected eye-related artifacts averaged across the cortex (top) and for significant frequency combinations (bottom). All coupling measures are z-scores relative to surrogate statistics generated by circular data shuffling. Opacity indicates statistical significance (p<0.05; corrected; permutation statistic).

### 3.2. Phase-amplitude coupling reflects residual muscle- and eye-related artifacts

Despite using ICA-based artifact cleaning and beamforming, the data may well contain residual artifacts. To investigate the potential influence of such residual artifacts, we computed vectorlength-PAC of MEG signals that mainly contained artifacts (Fig. 1B and 1C, Materials and Methods). If results reflected residual artifacts, we should observe similar patterns for artifact signals. Indeed, vectorlength-PAC of muscle artifacts (Fig. 1B) showed a prominent peak that well matched the high frequency peak in the cleaned data. The cortical distribution of vectorlength-PAC for muscle artifacts peaked in inferior temporal and frontal regions, the latter reminiscent of a saccadic spike artifact (Carl et al., 2012) (Fig. 1B). This cortical distribution was significantly correlated with the cortical distribution of PAC for the cleaned signal in the corresponding frequency range [*f*_*φ*_, *f*_*A*_] = [45-76Hz, 128-256Hz] (Fig. 1A, small inset) (r^2^ = 0.45, p<0.001). Together, these results suggested that high frequency PAC likely reflected muscle artifacts rather than neuronal coupling.

We repeated the same analysis for a potential contamination by residual eye-movement artifacts (Fig. 1C). Eye-movement artifacts showed low-frequency vectorlength-PAC for phase and amplitude frequencies below approximately 3 and 8 Hz, respectively. This only partially matched the weak but significant low frequency PAC for the cleaned data, which peaked for amplitude frequencies around 16 Hz reaching up to 64 Hz. Thus, low-frequency PAC could only be partially explained by residual eye-movement artifacts.

Next, we tested if these findings generalized to PAC estimation using bicoherence, which assess non-linear interactions between two frequencies by quantifying the phase consistency between these frequencies and their sum (Shahbazi Avarvand et al. 2018). Indeed, for high frequencies, we found a similar distribution of significant bicoherence for the cleaned data (Fig. 1D) and muscle artifacts (Fig. 1E). The cortical distribution of bicoherence of the cleaned data for this frequency range (Fig. 1D, left small inset) was significantly correlated with the cortical distribution of bicoherence for muscle-artifacts (Fig. 1E, left, r^2^=0.25, p<0.001) as well as with the cortical distribution of vectorlength-PAC of the cleaned data (Fig. 1A, small inset, r^2^=0.92, p<0.001) and muscle-artifacts (Fig. 1B, r^2^=0.46, p<0.001). This further suggested that residual muscle artifacts generated spurious high-frequency PAC in temporal, orbito-frontal and lateral inferior regions.

For eye movements artifacts, bicoherence could again not entirely explain PAC at low frequencies. Bicoherence of eye artifacts peaked at *f*_1_ frequencies from 1 to 2.5 Hz and *f*_2_ frequencies up to 20 Hz and above 30 Hz (Fig. 1F). In contrast, the cleaned data showed significant bicoherence over a broader low-frequency range (Fig. 1D). Moreover, the cortical distributions of bicoherence of eye-artifacts and the cleaned-data was only weakly correlated for low frequencies (r^2^=0.15, p<0.01).

In sum, both measures, vectorlength-PAC and bicoherence, consistently identified significant PAC that likely reflects residual muscle- and eye movement artifacts in high and low frequency ranges, respectively.

### 3.3. Phase-amplitude coupling in the alpha frequency-range

Vector-length PAC and bicoherence showed prominent effects in the alpha-frequency range that could not be attributed to muscle or eye movement artifacts. Do these effects reflect true neuronal interactions between distinct oscillations?

For vectorlength-PAC, the coupling of the alpha phase-frequency peaked at the double-phase-frequency diagonal (*f*_*A*_ = 2*f*_*φ*_), i.e. at the first higher harmonic of the phase frequency (maximum at [*f*_*φ*_, *f*_*A*_] = [11.3, 22.6]). Bicoherence showed several peaks at *f*_1_= 10.5 Hz and *f*_2_ of 11 Hz, 21 Hz, 30.5 Hz and 44 Hz, which are close to harmonics of *f*_1_. Thus, we hypothesized that these peaks may merely reflect harmonics of alpha oscillations, rather than true neuronal interactions. To address this question, we next focused on why PAC measures are sensitive to rhythmic signals with higher harmonics and how harmonics and non-harmonic PAC could be dissociated.

#### 3.3.1. Harmonic and non-harmonic phase-amplitude coupling

Figure 2 compares non-harmonic PAC, i.e. phase-amplitude coupling between two independent oscillations (Fig. 2 left, 10 Hz to 40 Hz coupling), with harmonic PAC, i.e. a rhythmic non-sinusoidal signal with higher harmonics (Fig. 2 right, 10 Hz base frequency with three harmonics).

**Figure 2.**
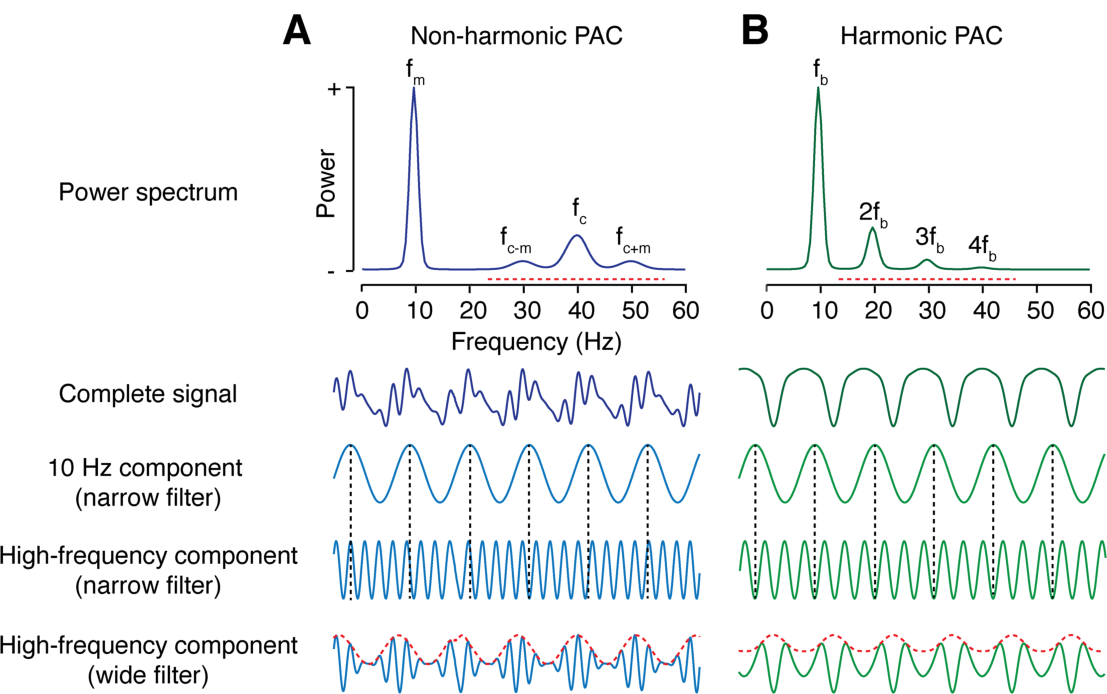
Non-harmonic and harmonic PAC. (**A**) Simulated signals with non-harmonic PAC between a modulating frequency *f*_*m*_ = 10 Hz and a carrier frequency *f*_*c*_ = 40 Hz (left) (**B** Simulated signal with harmonic PAC due to a non-sinusoidal signal with base frequency *f*_*b*_ = 10 Hz and three higher harmonics. First row: Power spectra. Red lines indicate the high-frequency bandwidth required for vectorlength-PAC. Second row: complete time-domain signals. Third row: low frequency component extracted with a narrow bandpass filter at 10 Hz. Fourth row: high frequency component of interest extracted with a narrow bandpass filter at *f*_*c*_ and 3*f*_*b*_ for non-harmonic and harmonic PAC, respectively. The narrow filter prevents amplitude modulation. For non-harmonic PAC, there is no stable phase relationship between the low and high frequency components. In contrast, for harmonic PAC, there is a stable phase relationship. Fifth row: high frequency components extracted with a wide filter (dashed red lines). Both signal types show an amplitude modulation of high-frequency components that is coherent with the 10 Hz component.

Importantly, both bicoherence and vectorlength-PAC do not only yield cross-frequency phase-amplitude coupling for non-harmonic PAC, but also for harmonic PAC (Kovach et al., 2018; Shahbazi Avarvand et al., 2018). How can this be intuitively understood?

For vectorlength-PAC, the spectral bandwidth of higher frequency components needs to be wide enough (at least twice the low frequency) to capture their potential amplitude modulation (Aru et al., 2015; Dvorak and Fenton, 2014, compare high frequencies extracted with narrow and wide filters in Fig. 2). For harmonic signals, this implies that multiple harmonics of the base frequency can be captured, which results in an apparent amplitude modulation of high frequencies (Fig. 2B). This modulation is phase-consistent with the base frequency, which yields significant vectorlength-PAC in absence of an independent high frequency oscillator.

Bicoherence does not necessitate wide filters but is also sensitive to both, harmonic and non-harmonic PAC (Avarvand et al. 2018). This is, because for harmonic PAC (Fig. 2, right), there is always a stable phase relationship between the base frequency and its higher harmonics. Thus, bicoherence measures cross-frequency coupling for all frequency pairs, where each frequency and their sum coincide with the base frequency or a higher harmonic.

#### 3.3.2. Dissociating harmonic and non-harmonic phase-amplitude coupling

How can we distinguish harmonic and non-harmonic PAC? Harmonics are only present at integer multiples of the base frequency. Thus, a first test is whether the frequencies at which PAC is observed could reflect harmonics of a base frequency. Bicoherence is particularly suited for this test, because it enables higher spectral resolution than vectorlength-PAC. However, if the observed PAC frequencies match a harmonic pattern, this may still reflect non-harmonic PAC at these frequencies (e.g. Fig. 2, left). Thus, for harmonic PAC frequency patterns, further tests are required.

To investigate such potential tests, we simulated signals that contained either harmonic or non-harmonic PAC between 10 Hz and some of its integer multiple frequencies. For each signal, we computed not only bicoherence to assess PAC, but also cross-frequency amplitude-amplitude coupling (AAC) and cross-frequency phase-phase coupling (PPC) (Fig. 3).

**Figure 3.**
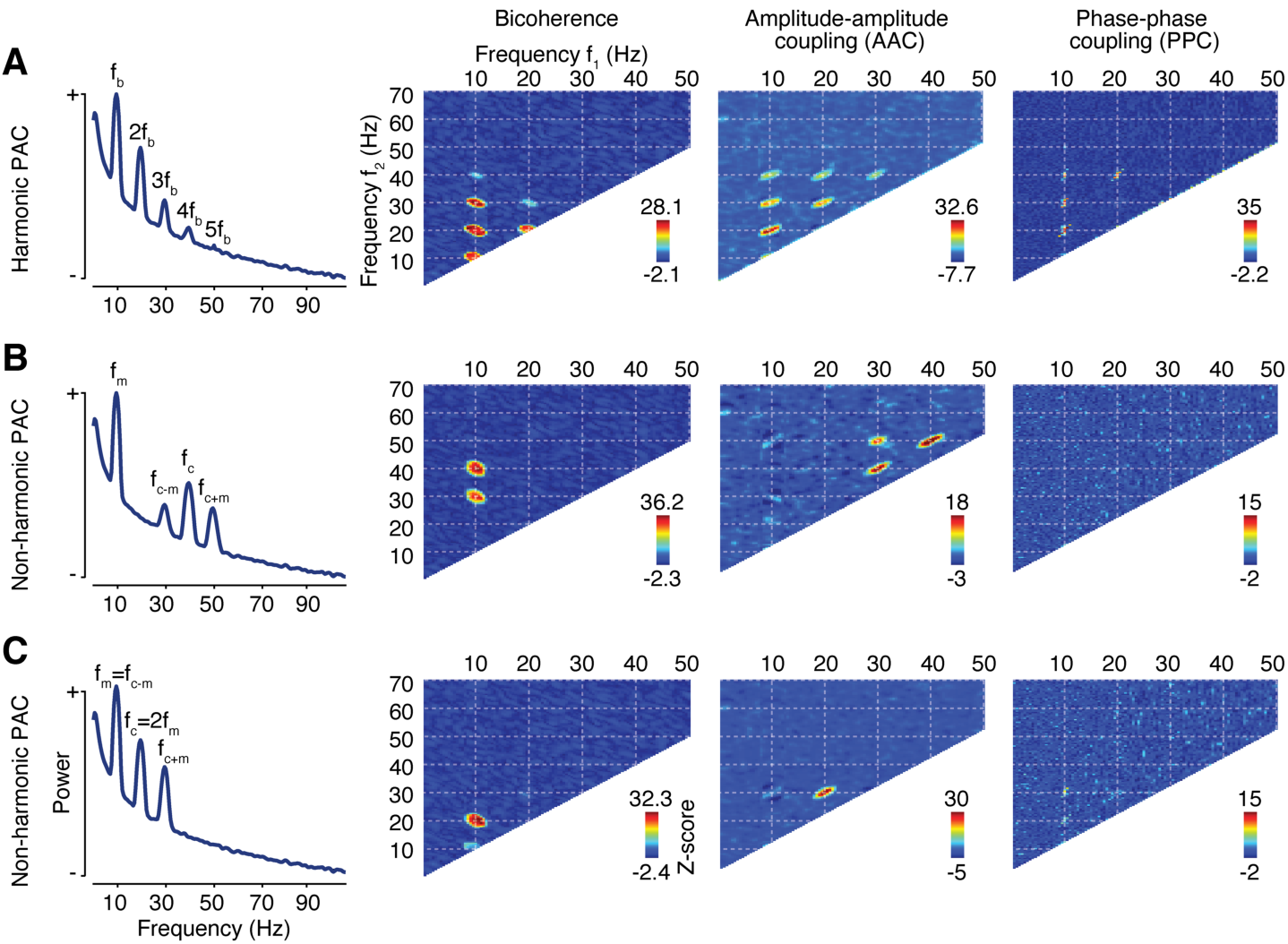
Dissociating harmonic and non-harmonic PAC. Power spectra, cross-frequency bicoherence, amplitude-amplitude coupling (AAC) and phase-phase coupling (PPC) for three different simulated signals with harmonic or non-harmonic PAC. All signals contained oscillatory components and uncorrelated 1/*f*^2^-noise. (**A**) Power and cross-frequency measures for a 10 Hz signal with four higher harmonics. (**B**) Non-harmonic PAC between 10 and 40 Hz. (**C**) Special case of non-harmonic PAC, between 10 Hz and 20 Hz. In this case, the lower side-peak of the modulated frequency *f*_*c-m*_ coincides with the modulating frequency *f*_*m*_. All coupling measures are z-scores relative to surrogate statistics generated by circular data shuffling.

For both, harmonic (Fig. 3A) and non-harmonic (Fig. 3B) PAC, bicoherence peaks at all [*f*_1_, *f*_2_] combinations, at which *f*_1_, *f*_2_, and *f*_3_ = *f*_1_ + *f*_2_ coincide with peaks in the power spectrum. For harmonic-PAC (Fig. 3A), power peaks at the base frequency and higher harmonics, which results in several bicoherence peaks. For non-harmonic PAC (Fig. 3B), power shows 4 peaks at the modulating frequency *f*_*m*_, carrier frequency *f*_*c*_ and side-peaks *f*_*m+/-c*_. This leads to two bicoherence peaks at frequencies [*f*_1_, *f*_2_] = [*f*_*m*_, *f*_*c*_] and [*f*_1_, *f*_2_] = [*f*_*m*_, *f*_*c-m*_] (Fig. 3B, Hyafil 2015). Thus, bicoherence with more than two peaks cannot only be caused by non-harmonic PAC and implies harmonic PAC.

Amplitude-amplitude and phase-phase coupling further dissociate harmonic and non-harmonic PAC. By definition, harmonics imply AAC and PPC between the base frequency and any harmonics. AAC and PPC measures of harmonic-PAC signals well reflect these couplings (Fig. 3A). In contrast, non-harmonic PAC does not imply AAC or PPC between the modulating frequency and the carrier frequency or its side-peaks. Thus, for non-harmonic PAC (Fig. 3B), there is no AAC or PPC between the modulating frequency and any of the power peaks at higher frequencies. This provides a dissociating feature between harmonic and non-harmonic PAC.

There is one special case of non-harmonic PAC, which is more ambiguous. This is when the carrier frequency is twice the modulating frequency (Fig. 3C). In this special case, the lower side-peak of the amplitude modulation coincides with the modulating frequency itself (10 Hz in Figure 3C). This leads to bicoherence and PPC patterns similar to patterns for a non-sinusoidal oscillator with only two higher harmonics. However, in this case, the relative strength of bicoherence and AAC peaks allows for dissociating harmonic and non-harmonic PAC, because the signal at the modulating frequency mixes with the lower-frequency side peak of the carrier frequency. For bicoherence, this leads to weaker coupling at [*f*_1_, *f*_2_] = [*f*_*m*_, *f*_*m*_] as compared to [*f*_1_, *f*_2_] = [*f*_*m*_, *f*_*c*_]. This is in contrast to harmonic PAC, which shows the opposite relative strength due to the lower power of harmonics as compared to the base frequency. Along the same line, for AAC coupling at [*f*_1_, *f*_2_] = [*f*_*m*_, *f*_*c*_] is absent or weaker than coupling at [*f*_1_, *f*_2_] = [*f*_*c*_, *f*_*c*_ + *f*_*m*_], which again is opposite to harmonic-PAC.

In sum, several features allow to assess if PAC is harmonic or non-harmonic in nature. A first useful heuristic is to test if the observed frequencies are multiples. If this is the case, bicoherence, AAC and PPC together are decisive. Bicoherence with more than two peaks implies harmonic PAC. Furthermore, bicoherence, AAC, and PPC between the base frequency and its harmonics imply harmonic PAC, in particular if bicoherence is strongest for identical *f*_1_ and *f*_2_ at the base-frequency. We next applied this approach to test if the observed PAC in the alpha range (Fig. 1A and 1D) reflected harmonic or non-harmonic PAC.

#### 3.3.3. Alpha phase-amplitude coupling reflects harmonics

As indicated above, average bicoherence peaked between alpha-frequencies *f*_1_ and harmonic frequencies *f*_2_ (Fig. 1B and 4A). This was also the case for every region with prominent PAC, i.e. sensorimotor, parietal, occipital and temporal areas (Fig. 4B–E). Therefore, we next computed AAC and PPC across the entire cortex and for the above four regions to assess the nature of alpha PAC (Fig. 4, middle and right column).

**Figure 4.**
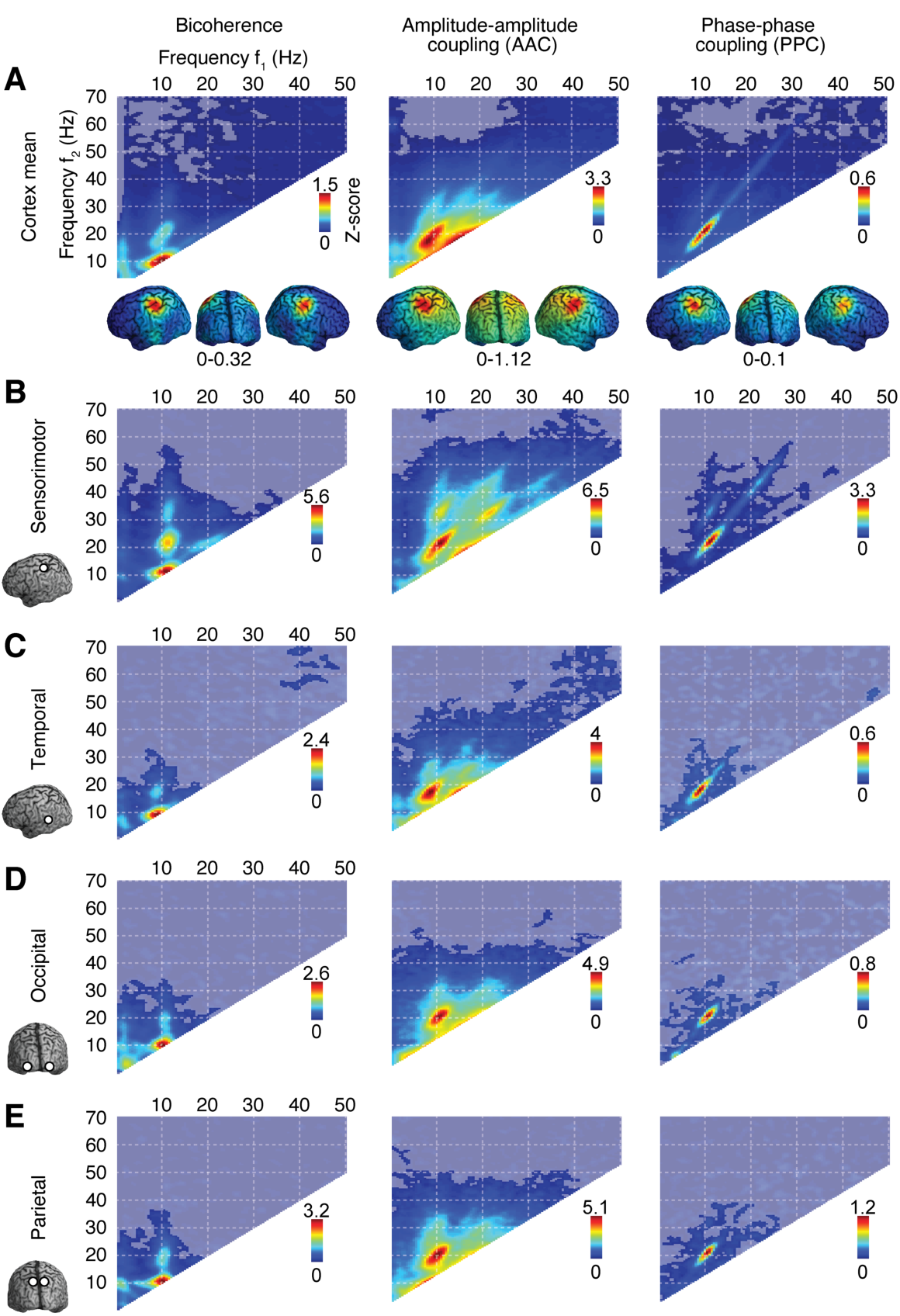
Cortical cross-frequency coupling. (**A**) Spectral and cortical distribution of bicoherence, cross-frequency amplitude-amplitude coupling and cross-frequency phase-phase coupling averaged across the cortex and across all frequency combinations, respectively. (**B-E**) Cross-frequency coupling measures for four different cortical ROIs. All coupling measures are z-scores relative to surrogate statistics generated by circular data shuffling. Opacity indicates statistical significance (p<0.05; corrected; permutation statistic).

The average across the cortex and all four regions of interest showed prominent AAC and PPC with strongest peaks at [*f*_1_, *f*_2_] = [10, 20] Hz as well as several effects at higher harmonics. Harmonic coupling was particularly strong for sensorimotor cortex, which showed prominent amplitude and phase coupling even between the second (20 Hz) and fourth (40 Hz) harmonic. We concluded that the observed PAC reflected harmonic-PAC due to non-sinusoidal alpha oscillations rather than non-harmonic PAC of independent oscillations.

If bicoherence, AAC and PPC all reflect the same harmonic PAC, their cortical distribution should be correlated. This is what we found. Averaged across all frequency pairs, the cortical patterns of bicoherence, AAC and PPC were strongly correlated (Bicoherence vs. AAC: r = 0.89, p < 0.0001; Bicoherence vs. PPC: r = 0.93, p < 0.0001; AAC vs. PPC: r = 0.88, p < 0.0001; Pearson correlation; Fig. 4A).

In sum, converging evidence suggested that PAC for low frequencies in alpha/beta range was harmonic in nature, and thus reflected non-sinusoidal signals.

#### 3.3.4. Harmonic phase-amplitude coupling reflects individual alpha rhythms

We next exploited the variability of the alpha-rhythm across subjects to further test the harmonic nature of alpha PAC. For harmonic-PAC the observed bicoherence peaks should scale with the variable alpha frequency across subjects. Thus, for each subject, we extracted all [*f*_1_, *f*_2_] peak frequencies for *f*_1_ in the alpha band and tested if these peaks were integer multiples of the individual alpha frequency (i.e. *f*_2_ = *n f*_1_, for *n* = 1,2,3).

For all regions, peak positions were compatible with the harmonic model (Fig. 5). The model fit of the lowest Bicoherence peak to the first harmonic at *f*_2_ = *f*_1_, which implies coupling between the base frequency and the second harmonic, was significant for all ROIs (motor ROI: r^2^ = 0.83, p<0.001; visual ROI: r^2^ =0.88, p<0.001; parietal ROI: r^2^ =0.92, p<0.001; temporal ROI: r^2^ =0.93, p<0.001). The model fit for the second harmonic (*f*_2_= 2*f*_1_) was significant for motor and temporal cortex (motor ROI: r^2^ = 0.85, p<0.001; temporal ROI: r^2^ = 0.88, p<0.05). For the motor cortex, also the model fit for the third harmonic (*f*_2_= 3*f*_1_) was significant (r^2^ = 0.87, p<0.001). Thus, for all regions of interest, the frequencies of bicoherence in individual subjects were well explained by alpha harmonics.

**Figure 5.**
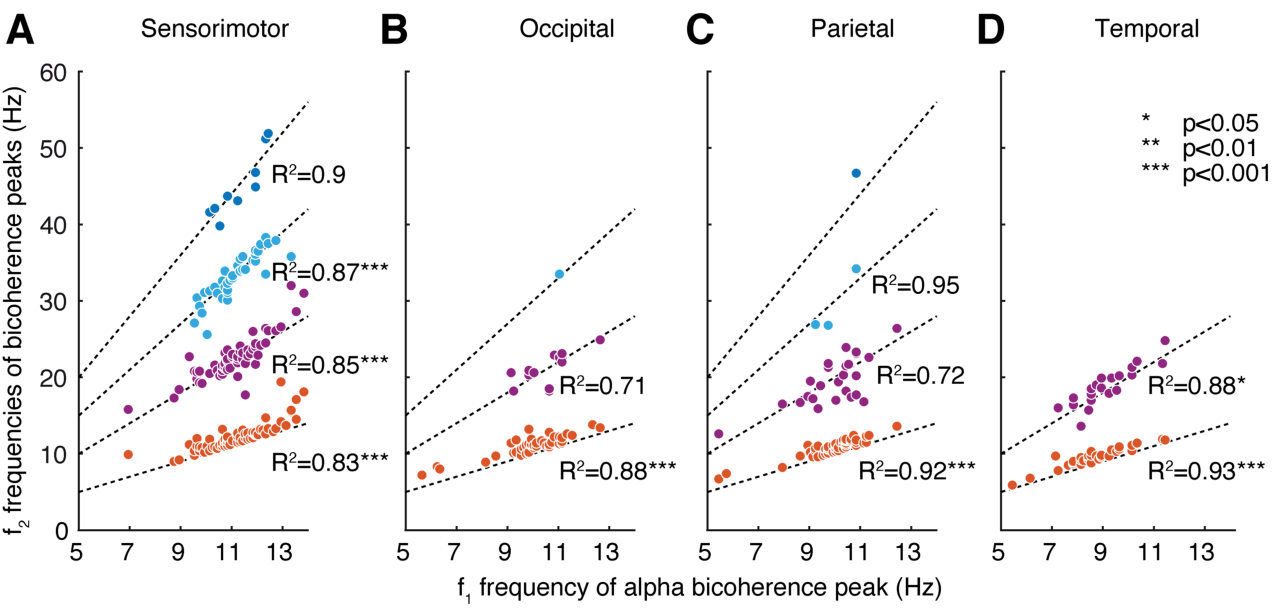
Harmonic PAC at individual alpha frequencies. (**A**) Bicoherence peak positions [*f*_1_, *f*_2_] with *f*_1_ frequencies in the alpha range across all individual subjects for all four ROIs. Dashed lines indicate harmonic models along with *R*^2^ values of model fits and statistical significances.

### 3.4. Replication in an independent dataset

We replicated the above findings in a second, independent dataset that was recorded with another MEG system at another research site (Fig. 6; Tübingen dataset). As for the HCP data, we found prominent bicoherence peaks at alpha *f*_1_ frequencies and at harmonic frequency combinations. Again, harmonic coupling was also present for AAC and PPC with decreasing coupling strengths for higher frequencies. We tested if the cortical distribution of PAC was similar between the two datasets. Indeed, the pattern of bicoherence, AAC and PPC was highly correlated between the two datasets (Bicoherence r=0.85, p<0.0001; AAC r = 0.85, p <0.0001; PPC r = 0.77, p<0.0001).

**Figure 6.**
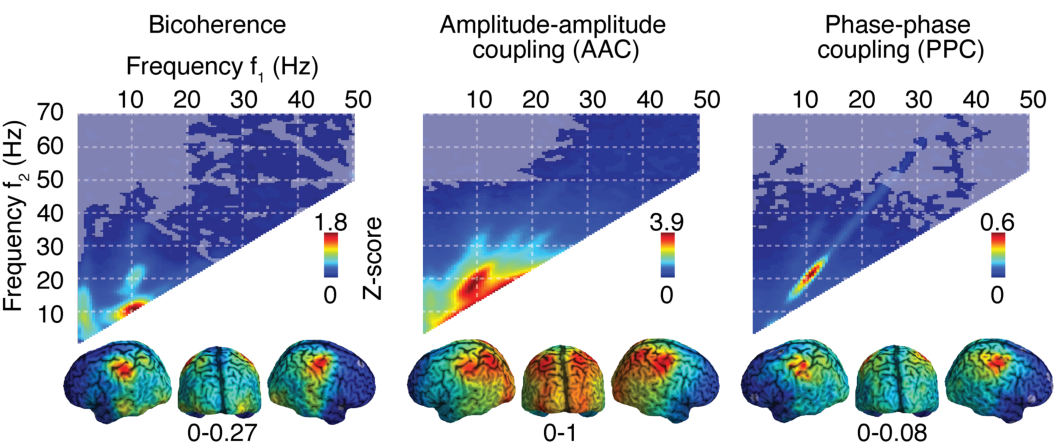
Replication of key results. (**A**) Spectral and cortical distribution of bicoherence, cross-frequency amplitude-amplitude coupling and cross-frequency phase-phase coupling averaged across the cortex and across all frequency combinations, respectively, for the replication dataset. All coupling measures are z-scores relative to surrogate statistics generated by circular data shuffling. Opacity indicates statistical significance (p<0.05; corrected; permutation statistic).

In sum, the results were highly consistent between the HCP and independent replication datasets. We concluded that, also for the replication dataset, PAC for low frequencies in the alpha range was harmonic in nature, and thus due to non-sinusoidal signals rather than due to non-harmonic PAC between independent oscillators.

### 3.5. Sub-alpha phase-amplitude coupling

Our initial vectorlength-PAC (Fig. 1A) and bicoherence analysis (Fig. 1D) revealed PAC at frequencies below 8 Hz that could not entirely be explained by eye-movement artifacts. In a final set of analyses, we investigated the nature of coupling in this sub-alpha frequency range.

The sensorimotor bicoherence of several subjects showed distinct peaks at sub-alpha *f*_1_ frequencies that seemed to mirror the harmonic peaks at the individual *f*_1_ alpha frequency (Fig. 7A). Thus, we hypothesized that these sub-alpha peaks might be caused by alpha-frequency harmonics. To investigate this, we analyzed low frequency bicoherence of a simulated 10 Hz signal with higher harmonics (Fig. 7B). Indeed, this revealed a characteristic “leakage pattern” at sub alpha frequencies with horizontal, vertical and anti-diagonal leakage lines that intersect at the harmonic 10 Hz peaks (*f*_2_=n*10, *f*_1_=n*10 and *f*_1_ + *f*_2_= n*10, respectively, for integer n). This specific pattern can be intuitively understood by the frequency combination used for bicoherence. Along horizontal, vertical or anti-diagonal lines, either the *f*_1_, *f*_2_ or *f*_2_ frequency stays constant. Consequently, the leakage of bicoherence is falling off slower along these lines than in any other direction.

**Figure 7.**
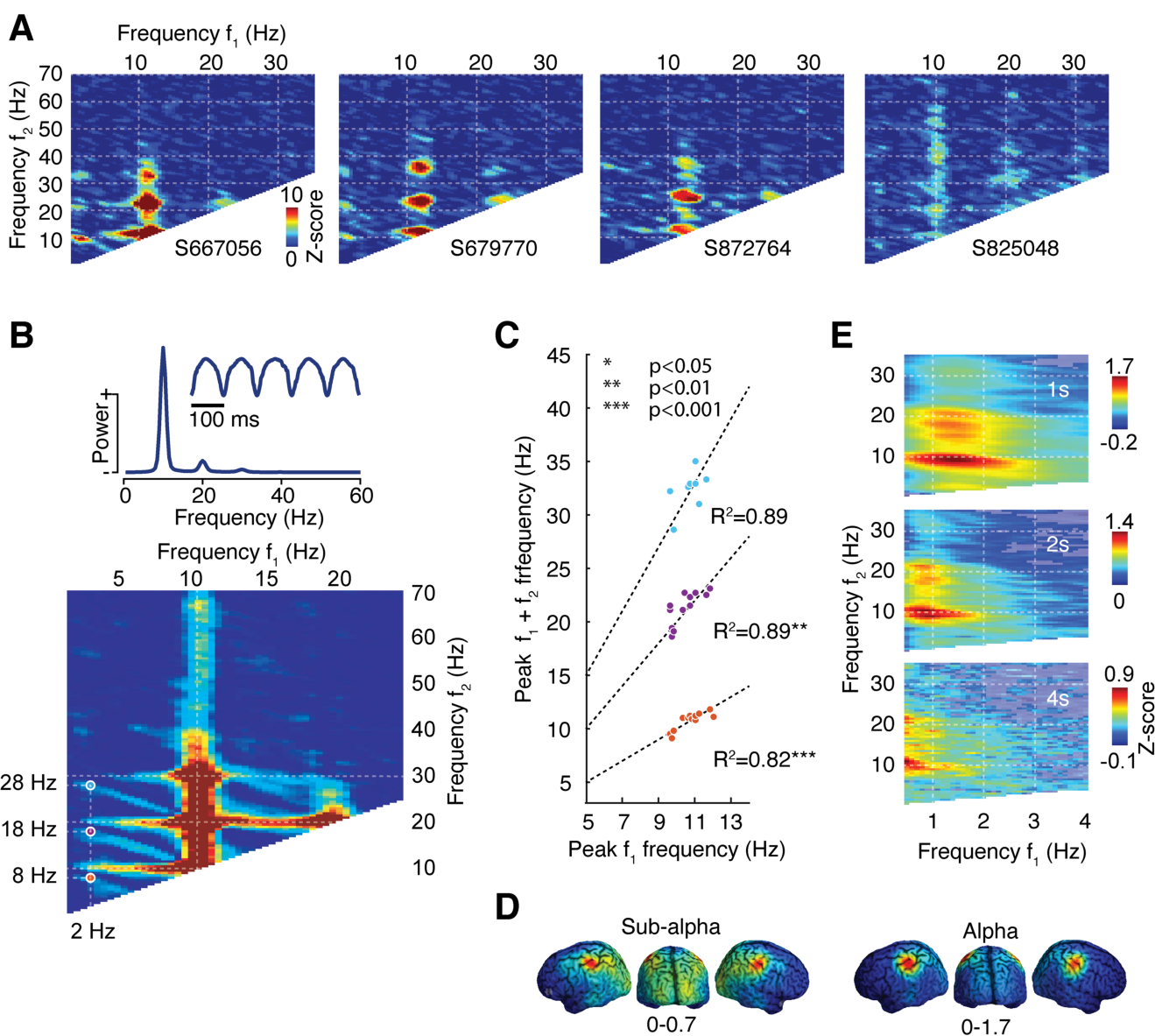
Spectral leakage causes spurious coupling. (**A**) Bicoherence from the motor ROIs of four subjects (**B**) top: simulated non-sinusoidal 10 Hz signal (power spectrum and time-domain); bottom: bicoherence of the simulated signal with apparent leakage-lines. Expected *f*_2_-leakage-postions corresponding to *f*_1_ = 2 Hz are marked. The sums of these *f*_1_ and *f*_2_ frequencies correspond to multiples of the base-frequency (10 Hz). (**C**) correspondence of *f*_1_ peak frequencies to the leakage model across subjects (**D**) similar cortical distribution of bicoherence in the sub-alpha and alpha range. (**E**) Sub-alpha bicoherence averaged across the cortex for 1s, 2s and 4s long temporal segmentation of the time-domain data (Hanning windows). All coupling measures are z-scores relative to surrogate statistics generated by circular data shuffling. Opacity in (E) indicates statistical significance (p<0.05; corrected; permutation statistic).

We next tested if such a leakage pattern was present in the MEG data. In each subject, we located up to three sub-alpha *f*_1_ bicoherence peaks and tested if their position was in line with the predicted leakage pattern (Fig. 7C). Indeed, peak positions were significantly correlated with the predictions of the leakage model across subjects (first peaks: R^2^=0.82, p <0.001; second peaks: R^2^ =0.89, p<0.01, third peaks R^2^ =0.89, p>0.05). This provided strong evidence that sub-alpha bicoherence peaks reflected spectral leakage of non-sinusoidal alpha oscillations. If this is correct, the cortical distribution of sub-alpha bicoherence should be correlated with the cortical distribution of alpha bicoherence. Indeed, we found a strong correlation of the cortical distribution of bicoherence in the leakage-related sub-alpha range ([*f*_1_, *f*_2_] = [0.5 to 4 Hz, 2.5 to 40 Hz]) and in the range of alpha harmonics ([*f*_1_, *f*_2_] = [7 to 14 Hz, 7 to 40 Hz] (r^2^ =0.81, p<0.001, Fig. 7D).

Notably, the above spectral analyses were based on temporal windowing using a 1 s Hanning taper. We hypothesized that the peak position of the leaked sub-alpha bicoherence was related to the spectral cut-off of this temporal windowing near 1.5 Hz. To test this, we repeated the analyses based on 2 s and 4 s Hanning windows (Fig. 7E). As hypothesized, the peak of leaked sub-alpha bicoherence shifted towards lower frequencies for longer temporal windows. Thus, the employed windowing modulates the exact spectral profile of PAC that is leaked below the frequency of non-sinusoidal oscillations

## 4. Discussion

We systematically investigated cross-frequency PAC across the human brain using MEG. We found no evidence for PAC that could be unequivocally attributed to non-harmonic coupling between distinct oscillators. Instead, consistent across two independent datasets, we observed cortically and spectrally wide-spread harmonic PAC reflecting non-sinusoidal oscillations in the alpha frequency-range.

### 4.1. Dissociating harmonic and non-harmonic PAC

Our findings accord well with previous reports that linked phase-amplitude coupling to alpha harmonics. Barnett et al. (1971) observed harmonic bicoherence, strongest over occipital and central areas, in subjects with strong alpha activity. Lozano-Soldevilla (2016) reported harmonic bicoherence of alpha ***f*_1_** frequencies in central and occipital MEG sensors. Following the increasing interest in cross-frequency interactions, the problem that PAC measures reflect harmonics has received growing attention (Aru et al., 2015; Chacko et al., 2018; Cole et al., 2017; Cole and Voytek, 2017; Gerber et al., 2016; Hyafil, 2015; Jensen et al., 2016; Kramer et al., 2008; Lozano-Soldevilla et al., 2016; Vaz et al., 2017; Velarde et al., 2019). Our findings well illustrate this problem and highlight the necessity to carefully distinguish between harmonic and non-harmonic PAC.

Our results show that the combination of three different cross-frequency coupling measures (bicoherence, amplitude coupling and phase coupling) allows not only to detect PAC, but also to reliably distinguish harmonic and non-harmonic PAC. Based on this approach, we characterized the nature of the observed cross-frequency interactions between alpha and higher frequencies. Our results identify harmonic-PAC, i.e. non-sinusoidal oscillations, as the mechanism underlying the abundant and strong PAC observed in this frequency range.

Our analytical approach well complements other methods or heuristics that can be used to dissociate harmonic and non-harmonic PAC. E.g. cross-frequency coupling could be related to neuronal spiking, PAC could be correlated to power, or PAC could be investigated between areas with distinct low- and fast oscillations (Jensen et al., 2016).

Here we focused on local PAC. Cross-area PAC (Chella et al., 2014; Nandi et al., 2019; Siebenhühner et al., 2020; van der Meij et al., 2012; von Nicolai et al., 2014) also needs to be carefully assessed with respect to harmonic signals. This is because local harmonics together with cross-area phase-coupling of the base-frequency may also result in measuring cross-area PAC without any underlying cross-area cross-frequency interaction between independent oscillations.

### 4.2. Physiological artifacts

Our results show that physiological artifacts such as eye-movements and muscle activity can cause spurious PAC and need to be carefully distinguished from brain activity. We observed strongest muscle artifacts in high frequency ranges reaching down to the alpha range, and eye-movement artifacts in the sub-alpha and sub-alpha to high-gamma range. Notably, we observed spurious PAC due to these artifacts although subjects fixated continuously, the data was cleaned for artifacts and source-reconstruction using beamforming further suppressed extracranial activity (Hipp and Siegel, 2013). This highlights the sensitivity of PAC measures to artifacts and the need to carefully exclude artifacts as causes of spurious PAC. This also includes other physiological artifacts such as heart-activity (Shahbazi Avarvand et al., 2018), which is typically projected to deeper sources.

### 4.3 Spectral leakage

Our results show that spectral leakage is an important caveat when assessing PAC. The simulation of a noisy non-sinusoidal alpha oscillation revealed a wide-spread pattern of spectral leakage of bicoherence between frequencies that were all linked to harmonics of alpha. We identified a matching leakage pattern in the MEG data suggesting that also sub-alpha PAC reflects leakage of harmonic alpha PAC. Critically, the leakage pattern was observed at lower frequencies than the base-frequency of the non-sinusoidal oscillator. Thus, non-sinusoidal signals can not only drive PAC at higher harmonics, but, due to spectral leakage, can also drive spurious PAC at lower frequencies than the base frequency at hand. This mechanism may be particularly problematic for muscle artifacts and may cause spurious PAC at very low frequencies typically not associated with muscle artifacts (compare Figs. 1E and 1F).

In this context, it should be noted that, while spectral leakage may be negligible for power, it may become particularly problematic for bicoherence and other cross-frequency measures. This is because, in contrast to power, these cross-frequency measures are sensitive to the preserved phase-consistencies between leaked signal components.

### 4.4. Spectral resolution

Our results show that alpha harmonics, leakage and residual muscle activity can impact broad frequency ranges. Thus, investigating only a few pre-defined frequency bands may lead to misinterpretations of PAC results. In other words, a comprehensive characterization of cross-frequency coupling across a broad frequency spectrum is required to unequivocally assess the nature of cross-frequency coupling. This in turn necessitates proper selection of spectral resolution.

Vectorlength-PAC requires a broad resolution (or wide filters) to detect phase-amplitude coupling (Aru et al., 2015). However, as our results show, this low resolution potentially masks informative spectral coupling patterns such as harmonic structures. Along the same line, also the use of a variable frequency resolution across the cross-frequency spectrum may be problematic, as it may mask informative spectral patterns. Bicoherence allows to avoid these problems and to obtain high-resolution PAC spectra (Shahbazi Avarvand et al. 2018). Our results show, how such bicoherence spectra can reveal harmonic coupling patterns and how they can be readily combined with cross-frequency amplitude- and phase-coupling spectra.

### 4.5. Open questions and conclusion

The fact that we did not find conclusive non-harmonic PAC in resting state MEG should be interpreted with caution. Our results may be specific to the resting-state. Non-harmonic PAC may be well observable in other task contexts. Furthermore, non-invasive population measures such as MEG may not reflect non-harmonic PAC present at the cellular or spiking level, which may be well detectable with invasive recordings. In any case, our findings show that PAC measures need to be interpreted with great care and taking into accounts several critical caveats highlighted here (Aru et al., 2015).

The prominent harmonic PAC shown here likely reflects the specific waveform of rhythmic processes, which, by itself, may be a highly informative physiological marker (Bartz et al., 2019; Cole et al., 2017; Cole and Voytek, 2017; Vaz et al., 2017). However, it should be noted that such harmonic PAC in principle may reflect both, a single non-sinusoidal rhythmic process as well as phase and amplitude coupling of multiple physiologically separate harmonic oscillations.

In sum, our results show how different cross-frequency measures can be combined to identify and characterize neuronal PAC. Using this approach, we did not observe conclusive non-harmonic PAC in human resting-state MEG. Instead, we found cortically and spectrally wide-spread harmonic PAC driven by non-sinusoidal alpha-band activity.

## CRediT authorship contribution statement

Janet Giehl: conceptualization, investigation, formal analysis, software, writing - original draft, review & editing. Nima Noury: conceptualization, writing - original draft, review & editing. Markus Siegel: conceptualization, writing - original draft, review & editing, resources, supervision, funding acquisition.

## Declarations of interest

none

## Acknowledgments

We thank Joerg Hipp and Anna Antonia Pape for recording part of the Tübingen MEG dataset. This study was supported by the European Research Council (ERC StG335880) and the Centre for Integrative Neuroscience (DFG, EXC 307). The authors declare no conflict of interest.

## Notes

### Competing Interest Statement

The authors have declared no competing interest.

## References

Aru, Juhan, Aru, Jaan, Priesemann, V., Wibral, M., Lana, L., Pipa, G., Singer, W., Vicente, R., 2015. Untangling cross-frequency coupling in neuroscience. Curr. Opin.Neurobiol. 31, 51–61. https://doi.org/10.1016/j.conb.2014.08.002

Barnett, T.P., Johnson, L.C., Naitoh, P., Hicks, N., Nute, C., 1971. Bispectrum analysis of electroencephalogram signals during waking and sleeping. Science 172, 401–402.

Bartz, S., Avarvand, F.S., Leicht, G., Nolte, G., 2019. Analyzing the waveshape of brain oscillations with bicoherence. NeuroImage 188, 145–160. https://doi.org/10.1016/j.neuroimage.2018.11.045

Benjamini, Y., Hochberg, Y., 1995. Controlling the False Discovery Rate: A Practical and Powerful Approach to Multiple Testing. J. R. Stat. Soc. Ser. B Methodol. 57, 289–300. https://doi.org/10.2307/2346101

Canolty, R.T., Edwards, E., Dalal, S.S., Soltani, M., Nagarajan, S.S., Kirsch, H.E., Berger, M.S., Barbaro, N.M., Knight, R.T., 2006. High gamma power is phase-locked to theta oscillations in human neocortex. Science 313, 1626–1628. https://doi.org/10.1126/science.1128115

Canolty, R.T., Knight, R.T., 2010. The functional role of cross-frequency coupling. Trends Cogn. Sci. 14, 506–515. https://doi.org/10.1016/j.tics.2010.09.001

Carl, C., Açık, A., König, P., Engel, A.K., Hipp, J.F., 2012. The saccadic spike artifact in MEG. NeuroImage 59, 1657–1667. https://doi.org/10.1016/j.neuroimage.2011.09.020

Chacko, R.V., Kim, B., Jung, S.W., Daitch, A.L., Roland, J.L., Metcalf, N.V., Corbetta, M., Shulman, G.L., Leuthardt, E.C., 2018. Distinct phase-amplitude couplings distinguish cognitive processes in human attention. NeuroImage 175, 111–121. https://doi.org/10.1016/j.neuroimage.2018.03.003

Chella, F., Marzetti, L., Pizzella, V., Zappasodi, F., Nolte, G., 2014. Third order spectral analysis robust to mixing artifacts for mapping cross-frequency interactions in EEG/MEG. NeuroImage 91, 146–161. https://doi.org/10.1016/j.neuroimage.2013.12.064

Cole, S.R., van der Meij, R., Peterson, E.J., de Hemptinne, C., Starr, P.A., Voytek, B., 2017. Nonsinusoidal Beta Oscillations Reflect Cortical Pathophysiology in Parkinson’s Disease. J. Neurosci. Off. J. Soc. Neurosci. 37, 4830–4840. https://doi.org/10.1523/JNEUROSCI.2208-16.2017

Cole, S.R., Voytek, B., 2017. Brain Oscillations and the Importance of Waveform Shape. Trends Cogn. Sci. 21, 137–149. https://doi.org/10.1016/j.tics.2016.12.008

Colgin, L.L., 2015. Theta–gamma coupling in the entorhinal–hippocampal system. Curr. Opin. Neurobiol. 31, 45–50. https://doi.org/10.1016/j.conb.2014.08.001

Dvorak, D., Fenton, A.A., 2014. Toward a proper estimation of phase–amplitude coupling in neural oscillations. J. Neurosci. Methods 225, 42–56. https://doi.org/10.1016/j.jneumeth.2014.01.002

Fell, J., Axmacher, N., 2011. The role of phase synchronization in memory processes. Nat. Rev. Neurosci. 12, 105–118. https://doi.org/10.1038/nrn2979

Gerber, E.M., Sadeh, B., Ward, A., Knight, R.T., Deouell, L.Y., 2016. Non-Sinusoidal Activity Can Produce Cross-Frequency Coupling in Cortical Signals in the Absence of Functional Interaction between Neural Sources. PloS One 11, e0167351. https://doi.org/10.1371/journal.pone.0167351

Gross, J., Kujala, J., Hamalainen, M., Timmermann, L., Schnitzler, A., Salmelin, R., 2001. Dynamic imaging of coherent sources: Studying neural interactions in the human brain. Proc. Natl. Acad. Sci. U. S. A. 98, 694–699. https://doi.org/10.1073/pnas.98.2.694

Hagihira, S., Takashina, M., Mori, T., Mashimo, T., Yoshiya, I., 2001. Practical issues in bispectral analysis of electroencephalographic signals. Anesth. Analg. 93, 966–970, table of contents.

Hipp, J.F., Siegel, M., 2015. BOLD fMRI Correlation Reflects Frequency-Specific Neuronal Correlation. Curr. Biol. CB 25, 1368–1374. https://doi.org/10.1016/j.cub.2015.03.049

Hipp, J.F., Siegel, M., 2013. Dissociating neuronal gamma-band activity from cranial and ocular muscle activity in EEG. Front. Hum. Neurosci. 7, 338. https://doi.org/10.3389/fnhum.2013.00338

Hyafil, A., 2015. Misidentifications of specific forms of cross-frequency coupling: three warnings. Front. Neurosci. 9, 370. https://doi.org/10.3389/fnins.2015.00370

Hyafil, A., Giraud, A.-L., Fontolan, L., Gutkin, B., 2015. Neural Cross-Frequency Coupling: Connecting Architectures, Mechanisms, and Functions. Trends Neurosci. 38, 725–740. https://doi.org/10.1016/j.tins.2015.09.001

Jensen, O., Bonnefond, M., VanRullen, R., 2012. An oscillatory mechanism for prioritizing salient unattended stimuli. Trends Cogn. Sci. 16, 200–206. https://doi.org/10.1016/j.tics.2012.03.002

Jensen, O., Colgin, L.L., 2007. Cross-frequency coupling between neuronal oscillations. Trends Cogn. Sci. 11, 267–269. https://doi.org/10.1016/j.tics.2007.05.003

Jensen, O., Spaak, E., Park, H., 2016. Discriminating Valid from Spurious Indices of Phase-Amplitude Coupling. eNeuro 3. https://doi.org/10.1523/ENEURO.0334-16.2016

Kovach, C.K., Oya, H., Kawasaki, H., 2018. The bispectrum and its relationship to phase-amplitude coupling. NeuroImage 173, 518–539. https://doi.org/10.1016/j.neuroimage.2018.02.033

Kramer, M.A., Tort, A.B.L., Kopell, N.J., 2008. Sharp edge artifacts and spurious coupling in EEG frequency comodulation measures. J. Neurosci. Methods 170, 352–357. https://doi.org/10.1016/j.jneumeth.2008.01.020

Lisman, J., Idiart, M., 1995. Storage of 7 +/- 2 short-term memories in oscillatory subcycles. Science 267, 1512–1515. https://doi.org/10.1126/science.7878473

Lozano-Soldevilla, D., Ter Huurne, N., Oostenveld, R., 2016. Neuronal Oscillations with Non-sinusoidal Morphology Produce Spurious Phase-to-Amplitude Coupling and Directionality. Front. Comput. Neurosci. 10, 87. https://doi.org/10.3389/fncom.2016.00087

McLelland, D., VanRullen, R., 2016. Theta-Gamma Coding Meets Communication-through-Coherence: Neuronal Oscillatory Multiplexing Theories Reconciled. PLOS Comput. Biol. 12, e1005162. https://doi.org/10.1371/journal.pcbi.1005162

Nandi, B., Swiatek, P., Kocsis, B., Ding, M., 2019. Inferring the direction of rhythmic neural transmission via inter-regional phase-amplitude coupling (ir-PAC). Sci. Rep. 9, 6933. https://doi.org/10.1038/s41598-019-43272-w

Oostenveld, R., Fries, P., Maris, E., Schoffelen, J.-M., 2011. FieldTrip: Open source software for advanced analysis of MEG, EEG, and invasive electrophysiological data. Comput. Intell. Neurosci. 2011, 156869. https://doi.org/10.1155/2011/156869

Shahbazi Avarvand, F., Bartz, S., Andreou, C., Samek, W., Leicht, G., Mulert, C., Engel, A.K., Nolte, G., 2018. Localizing bicoherence from EEG and MEG. NeuroImage 174, 352–363. https://doi.org/10.1016/j.neuroimage.2018.01.044

Siebenhühner, F., Wang, S.H., Arnulfo, G., Lampinen, A., Nobili, L., Palva, J.M., Palva, S., 2020. Genuine cross-frequency coupling networks in human resting-state electrophysiological recordings. PLOS Biol. 18, e3000685. https://doi.org/10.1371/journal.pbio.3000685

Siegel, M., Donner, T.H., Engel, A.K., 2012. Spectral fingerprints of large-scale neuronal interactions. Nat. Rev. Neurosci. 13, 121–134. https://doi.org/10.1038/nrn3137

Tort, A.B.L., Komorowski, R., Eichenbaum, H., Kopell, N., 2010. Measuring phase-amplitude coupling between neuronal oscillations of different frequencies. J. Neurophysiol. 104, 1195–1210. https://doi.org/10.1152/jn.00106.2010

van der Meij, R., Kahana, M., Maris, E., 2012. Phase-Amplitude Coupling in Human Electrocorticography Is Spatially Distributed and Phase Diverse. J. Neurosci. 32, 111–123. https://doi.org/10.1523/JNEUROSCI.4816-11.2012

Van Essen, D.C., Smith, S.M., Barch, D.M., Behrens, T.E.J., Yacoub, E., Ugurbil, K., WU-Minn HCP Consortium, 2013. The WU-Minn Human Connectome Project: an overview. NeuroImage 80, 62–79. https://doi.org/10.1016/j.neuroimage.2013.05.041

Van Veen, B.D., van Drongelen, W., Yuchtman, M., Suzuki, A., 1997. Localization of brain electrical activity via linearly constrained minimum variance spatial filtering. IEEE Trans. Biomed. Eng. 44, 867–880. https://doi.org/10.1109/10.623056

Vaz, A.P., Yaffe, R.B., Wittig, J.H., Inati, S.K., Zaghloul, K.A., 2017. Dual origins of measured phase-amplitude coupling reveal distinct neural mechanisms underlying episodic memory in the human cortex. NeuroImage 148, 148–159. https://doi.org/10.1016/j.neuroimage.2017.01.001

Velarde, O.M., Urdapilleta, E., Mato, G., Dellavale, D., 2019. Bifurcation structure determines different phase-amplitude coupling patterns in the activity of biologically plausible neural networks. NeuroImage 202, 116031. https://doi.org/10.1016/j.neuroimage.2019.116031

von Nicolai, C., Engler, G., Sharott, A., Engel, A.K., Moll, C.K., Siegel, M., 2014. Corticostriatal coordination through coherent phase-amplitude coupling. J. Neurosci. Off. J. Soc. Neurosci. 34, 5938–5948. https://doi.org/10.1523/JNEUROSCI.5007-13.2014

